# Deciphering the Inter-domain Coupling in a Gram-negative Bacterial Membrane Insertase

**DOI:** 10.1101/2022.08.09.503346

**Authors:** Adithya Polasa, Shadi A Badiee, Mahmoud Moradi

## Abstract

YidC is a membrane protein that plays an important role in inserting newly generated proteins into lipid membranes. The Sec-dependent complex is responsible for inserting proteins into the lipid bilayer, and this process is facilitated by YidC in bacteria. In addition, YidC acts as a chaperone during the folding process of proteins. Multiple investigations have conclusively shown that the gram-positive bacterial YidC has Sec-independent insertion mechanisms. Through the use of microsecond-level allatom molecular dynamics (MD) simulations, we have carried out the first in-depth investigation of the YidC protein originating from gram-negative bacteria. This research sheds light on the significance of multiple domains of YidC structure at an atomic level by utilizing equilibrium MD simulations. Specifically, in this research, multiple models of YidC embedded in the lipid bilayer were constructed to characterize the critical role of the C2 loop and the periplasmic domain present in gram-negative YidC, which is absent in its gram-positive counterpart. Based on our results, the C2 loop is responsible for the overall stabilization of the protein, most notably in the transmembrane region, and it also has an allosteric influence on the periplasmic domain. We have found critical inter- and intra-domain interactions that contribute to the stability of the protein and its function. Finally, our study provides a hypothetical Sec-independent insertion mechanism for gram-negative bacterial YidC.

## Introduction

Membrane proteins participate in fundamentally crucial biological processes such as signaling, transcriptional regulation, ion transport, proteolysis, motility, metabolism, energy creation, and energy transfer^1^. In bacteria certain specialized cellular machineries enable proper membrane protein folding and insertion into the lipid bilayer^2–4^. Many membrane proteins are inserted through the Sec apparatus. YidC is one of the proteins working with Sec machinery in order to introduce client proteins into the membrane^2–8^.

YidC is a member of the Oxa/Alb3/YidC family of insertases found in mitochondria, chloroplasts, and bacteria^9–15^. YidC catalyzes the transmembrane insertion of freshly produced membrane proteins in the absence of an energy supply domain, such as an ATPase^16^. It also plays a vital role in the insertion and positioning of membrane proteins in bacteria^17–19^. Insertase proteins, such as YidC, have been exhaustively investigated to determine their importance for the insertion of proteins into membranes^20^, and many researchers have found evidence of YidC in conjunction with the Sec-complex^2–8^, which act to insert peptides into the membrane bilayer through the Signal Recognition Particle (SRP) mechanism. In addition, YidC may fold and insert polypeptides independent of the Sec-dependent pathway,^3,21–28^ and it is essential for the insertion of small phage coat proteins like Pf3 coat and M13 in a Sec-independent pathway.^21,29–34^.

A few experimental studies have explored the role of YidC in various microbial organisms^35–40^. The genomes of the majority of gram-positive microscopic organisms encode the YidC1 and YidC2 proteins.^38,39^. Although YidC typically exists as a dimer or tetramer^41,42^ under physiological conditions, it is discovered that YidC can also exist as a monomer in lipid bilayers^16,43^. On the other hand, the gram-negative YidC protein possesses an additional transmembrane (TM) segment at the N-terminus and a large periplasmic domain (PD)^44,45^. Although the PD region and the additional N-terminus TM segment are not required for YidC functional activity, the PD region interacts with Sec machinery and helps to create a stable complex^44^. The area with the C-terminal five TM segments is vital for the membrane insertase activity of gram-negative YidC^37^. In both gram-negative and gram-positive bacterial strains, the protein is firmly anchored within the lipid bilayer by interfacial aromatic residues, a cytoplasmic salt-bridge group, and a periplasmic helix enhanced with aromatic residues^35–38^. The highly conserved arginine residue (R366) in the hydrophilic groove was found in the same locations as in gram-positive (R72), implying that the arginine residue is as important for the function of gram-negative bacteria as it is for gram-positive bacteria^44^. The C-terminus of monomeric YidC interacts with the ribosomes, and the short interhelical loops C1 and C2 come into contact with the ribosomal proteins^46^. A group of aromatic residues around R72/R366 may bind with incoming peptide during insertion into the lipid bilayer^20,35–40^.

YidC is hypothesized to facilitate membrane insertion in both gram-negative and grampositive bacteria by binding incoming peptides through cytoplasmic loop interactions, hydrophobic force, and groove interactions^16,20,47,48^. The hydrophilic groove inside the membrane core of YidC increases the rate of accepting the hydrophilic moieties of a substrate into the membrane^49–52^. During its independent insertion mechanism, gram-positive YidC also goes through several conformational changes, including widening of the TM region and hydration and dehydration of the hydrophilic groove^20^. Additionally, a broad range of interactions with the incoming protein is involved at each step of the insertion process. For instance, the salt bridge interaction with R72 aromatic residue^20^.

While the YidC insertase in gram-positive bacteria has been extensively investigated in previous studies^16,20,47,48^, the significance of the additional periplasmic domain (PD) in gram-negative YidC still needs to be understood. There is insufficient evidence to determine whether the PD region of YidC influences protein stability and function. Moreover, whether the cytoplasmic loops play analogous roles in gram-positive and gram-negative bacterial YidC is unclear. Here, gram-negative YidC’s structure was investigated using microsecond-level all-atom MD simulations. We examined the local and global conformational changes in YidC brought on by the loss of the PD and the cytoplasmic C2 loop.

## Methods

From the Protein Data Bank, the gram-negative bacterial YidC crystal structure (PDB:6al2^53^) was downloaded. The CHARMM36m^54^ force field^55^, together with the NAMD 2.14^56^ software package was used for all-atom Molecular Dynamics (MD) simulations. Using the membrane builder on CHARMM-GUI^57^, YidC was introduced into the lipid bilayer, solvated, and ionized. In these MD investigations, YidC was embedded in a lipid bilayer of 1-palmitoyl-2- oleoyl-sn-glycero-3-phosphoethanolamine (POPC) lipids. A 90 Å × 90 Å membrane layer surface was constructed along the XY plane. The protein-lipid assembly was solvated in TIP3^58^ water with 18 Å thick layers of water on top and bottom. To neutralize the system, 0.15 M of Na^+^ and Cl*^−^* ions were added to the solution, with a slight modification in the number of ions. There were about *≈*143000 atoms in the final solvated system. Utilizing the conjugate gradient technique^59^, each system underwent energy minimization prior to the equilibrium simulation. The systems were then gradually relaxed using constrained MD simulations in accordance with the standard CHARMM-GUI^57^ procedure. In the NPT ensemble at 310 K, 1 *µ*s of equilibrium MD simulations were performed under periodic boundary conditions for each system. In the simulations, a Langevin integrator with a damping coefficient of *γ* = 0.5 ps*^−^*^1^ and 1 atm pressure was maintained using the Nose-Hoover Langevin piston method^19,60–71^.

We are interested in the significance of the cytoplasmic C2 loop and extracellular periplasmic domain (PD). Previous studies have highlighted the critical role of the C2 loop in determining YidC’s conformation and function in gram-positive bacteria^20,72^. We are interested in learning more about the functions of the PD (Fig. 1A), which is absent in gram-positive bacterial YidC and present in gram-negative bacterial YidC. Accordingly, we created four systems: YidC with PD and C2 loop (YidC), YidC without C2 loop (YidC ΔC2), YidC without PD region (YidC ΔPD), and YidC without PD region and C2 loop (YidC ΔPD ΔC2).

**Fig. 1.**
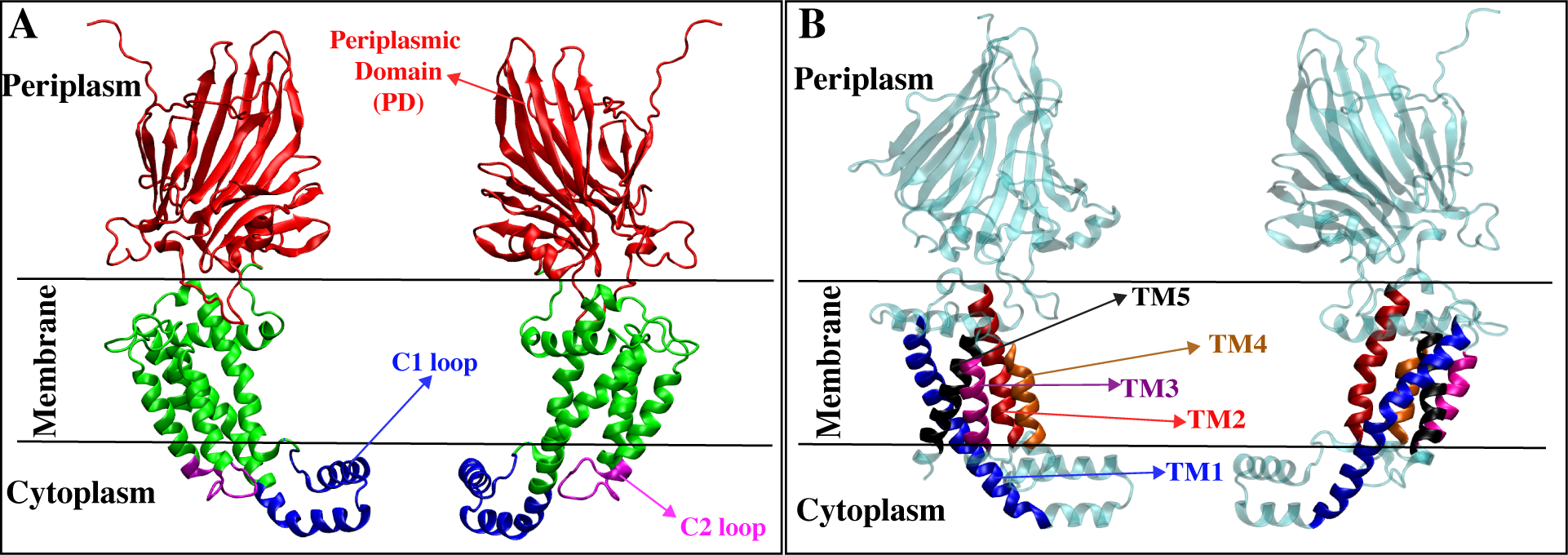
The cartoon representation of YidC (PDB:6AL2) (A) includes periplasmic domain (PD), transmembrane (TM) domain, C1 and C2 loops on the cytoplasmic side. (B) The cartoon representation of YidC’s individual TM helices TM1 (blue), TM2 (Red), TM3 (purple), TM4 (orange), and TM5 (black).

The preliminary simulations were run on TACC Stampede supercomputer. Subsequently, the production run for each model was extended to 400 ns with a timestep of 2.5 fs on Anton2^62^. Every 240 picoseconds, conformations were gathered on Anton2. The Anton2 simulation trajectories were initially processed on Kollman^62^. Later, the simulations were extended to 1 microsecond on TACC Stampede, and one additional set of 1 *µ*s simulation was performed for all systems.

All trajectories were visualized and examined using the VMD software^73^. A VMD plugin was used to analyze salt bridge interactions by measuring the distance between the oxygen atoms of acidic residues and the nitrogen atoms of basic residues with a cut-off distance of 4 Å. The interhelical angles were determined as the angle between the third main axes of the respective helices^20,64,71,74,75^. The residue selection of the TM helices and other sub-domains are as indicated: TM1 (355-388); TM2 (423-442); TM3 (466-479); TM4 (497-508); TM5 (511-528); C1 loop (380-420); C2 loop (480-492); and PD (49-326) (Fig. 1). To analyze the water inside the groove region, we counted the number of water molecules within 5 Å of R366. Principle component analysis (PCA) was performed for each trajectory using PRODY^20,64,76^, taking only protein *C_α_* atoms into account.

More quantitative data on the coordinated movements of the *C_α_* atoms was obtained using Dynamic Network Analysis (DNA) of the associated motions of the protein^62,77^. The correlation coefficient for the motion of each *C_α_* atom with respect to the other *C_α_* atoms was determined using MD-TASK^78^, a software package of MD analysis tools. For each of the TM regions in all the simulated trajectories, a correlation matrix *M* was created.

To quantify the differences in correlation between a system and some reference, a difference matrix Δ was calculated,

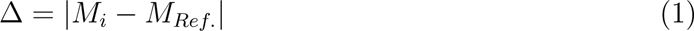

where *M_i_* is the correlation matrix of interest, and *M_Ref_* is the correlation matrix of a reference conformation. In this work, our point of interest was the difference between a TM region in complete YidC structure conformation and other YidC structures. For this reason, the TM region in the wild-type YidC simulations was compared with the TM region in the other YidC simulation system described above.

## Results and Discussion

### The Overall Protein Conformation is Stabilized by the Presence of the C2 loop and the PD Domain

The protein’s root mean square deviation (RMSD) was first calculated to assess its stability during equilibrium simulations. The RMSD of the YidC TM region in respect to the initial frame was computed, and the results are shown as a function of the simulation run-time (Fig. 2). According to the C*_α_* RMSD of the four different systems, the presence of the PD and C2 loop helps to stabilize the protein when it is in its native state (Fig. 2A). The system without the C2 loop is, beyond a shadow of a doubt, less stable than the system with the loop (Fig. 2B & D). The combined effect of removing the PD and the C2 loop has a higher impact on RMSD (Fig. 2D), which suggests that the PD could be responsible for preserving the stability of the protein. However, the effect of just removing the C2 loop (Fig. 2B) on protein RMSD is slightly higher than the native system (Fig. S1), but not as high as the system without PD and the C2 loop (Fig. 2D).

**Fig. 2.**
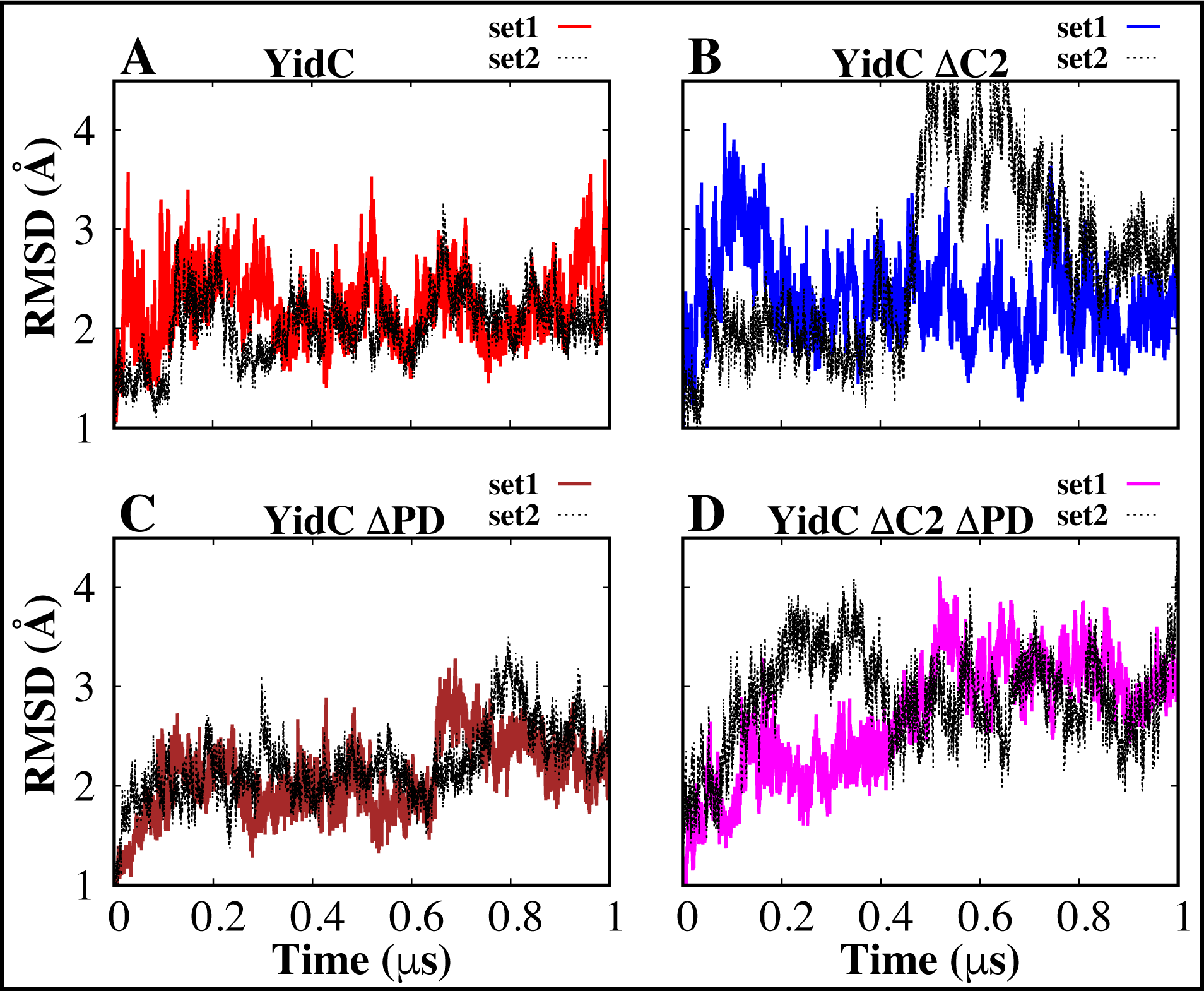
Analysis of YidC’s structural stability in the presence and absence of PD and C2 loop. (A–D) The root mean square deviation of the YidC in different systems. Based on RMSD data, we have observed that YidC fluctuates more in the system with the C2 loop removed compared to systems where the C2 loop is present. The simulations for each system were run twice, and the dashed lines in the graphs reflect the second run of those simulations.

This indicates the C2 loop is more important for protein stability than the PD region, as evidenced in the system YidC ΔPD (Fig. 2C). The RMSD of the TM region is very similar to the wild-type YidC structure (Fig. 2A).

Therefore, we have concluded that the impact of eliminating the C2 loop on gram-negative bacterial YidC is far more substantial than the effect of removing just PD regions (Fig. S1). The principal component analysis (PCA) was employed to identify the most significant differences between the systems. The projections onto principal components (PCs) 1 and 2 facilitated a clear distinction between the YidC wild type and the other systems (Fig. 3).

**Fig. 3.**
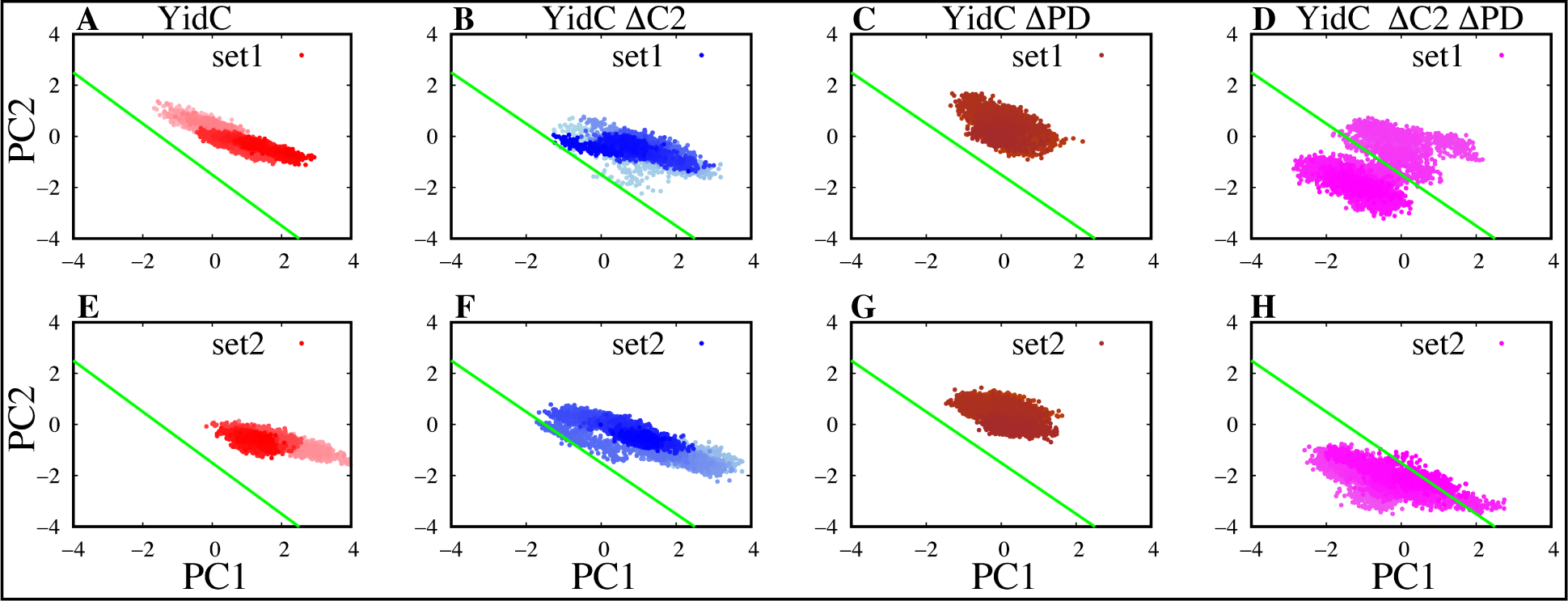
Projections of the principal components (PCs) 1 and 2. (A–H) PCA findings of PC1 versus PC2 for YidC systems from set 1 simulations are shown in the top row, while the results of set 2 simulations are displayed in the bottom row. The gradation of colors in the image indicates a timeline, with lighter shades reflecting earlier points in the simulation and darker color indicating later points. For consistency, only the PCA analysis of the transmembrane (TM) area, present in all systems, is displayed here. To facilitate comparison, the green line on the plot indicates the difference in PC projections.

In this particular analysis, only the YidC *C_α_* atoms located in the TM area are considered. PC1 contributed 27.5% of the total variance, while PC2 contributed 19.1% of the total variance. It was anticipated that the structural analysis of YidC ΔC2 and YidC ΔC2 ΔPD models would contradict that of YidC and YidC ΔPD in PC1 and PC2 (Fig. 3). This contradiction is rational, considering the significant conformational discrepancies found earlier in the RMSD analysis (Fig. S1). In general, the most important thing that came out of the principal components analysis was the realization that the behavior of the YidC ΔC2 ΔPD (Fig. 3 D & H) protein system was quite different from the other systems (Fig. 3). The results of this study support the theory that the C2 loop plays a significant part in the conformational dynamics of YidC.

Previous studies have revealed that the YidC transmembrane (TM) region is crucial for the membrane protein insertion mechanism^20,79,80^. In order to examine the impact of deleting the PD and C2 loop on the TM helices, the helical angle between each pair of TMs was measured in this work (Fig. 4). Compared to the wild type, we found that the local shape of the TM helices was altered in all of the other systems. Similar to the analysis above, the local conformation was more affected by the removal of both the PD and the C2 loop (Fig. 4D & H). However, when just the PD is removed, we do see some changes in the helical angle (Fig. 4C & G), despite the fact that the impact is not as significant as what is seen in systems that do not have a C2 loop (Fig. 4B, F, D, & H). This demonstrates the critical role that the C2 loop plays in maintaining the structural stability of the transmembrane region of YidC. We do observe a similar trend in other combinations of transmembrane helices (Fig. S2).

**Fig. 4.**
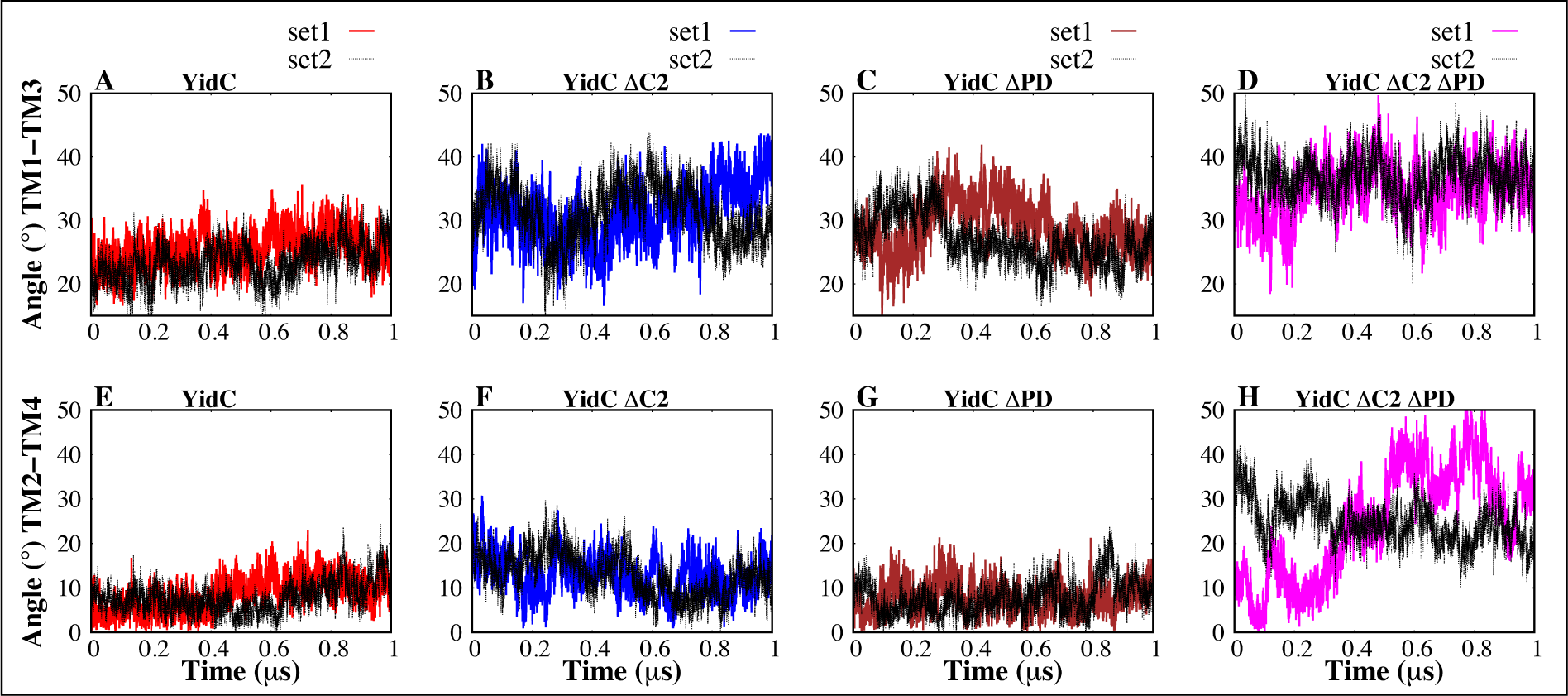
Inter-helical angles between transmembrane helices of YidC. (A–D) The interhelical angle between the protein’s transmembrane helix 1 and helix 3. (E–H) The overall inter-helical angle between helix 2 and helix 4 of the protein.

Furthermore, we used dynamic network analysis (DNA), which finds the linear connection between various residue pairs, to conduct a comprehensive study on the allosteric interactions of the C2 loop and the PD region with the various protein domains. The correlation coefficient of each residue pair is shown in Figure 5. This coefficient was calculated from the trajectory with the C2 loop and PD region, then subtracted from the same quantity calculated from the trajectory without the loop and PD region systems, and the result was reported as its absolute value. The amount presented for each pair of residues measures the size of the difference in the correlation behavior of the two residues brought on by the presence of the C2 loop and the PD domain in the TM region. The presence or absence of the C2 loop has been demonstrated to create significant variations in the correlations between YidC’s distinct domains. The inter-domain correlations, notably between the TM/C1 loop region, vary significantly between the YidC and other systems (Fig. S3). This difference in the cross-correlation is more pronounced, especially between TM1 and TM4 (Fig. 5). Based on this, we believe that the C2 loop does play a crucial role in the YidC conformational dynamics and that the conformational dynamics of the TM region are affected in its absence. The DNA findings (Fig. S3) are consistent with the early evidence for global and regional structural alterations. Overall, the results show that the C2 loop affects the behavior of the functionally essential areas of gram-negative YidC.

**Fig. 5.**
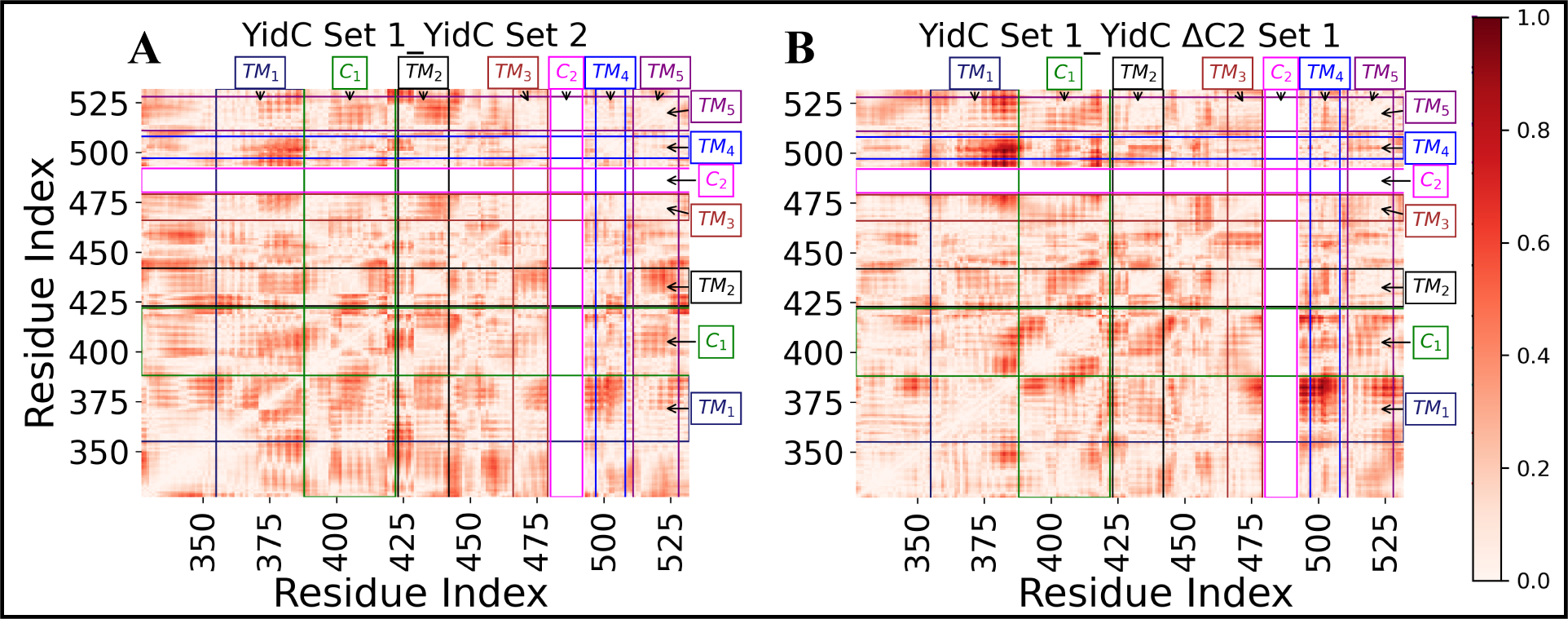
The DNA analysis revealed differences in correlation between the YidC set 1 control system and the YidC ΔC2 ΔPD system examined. The theoretical maximum for correlation difference is 2, but the observed maximum was less than 1. (A–B) Differences in correlation are shown as a red gradient, with darker red indicating a more significant difference.

### C2 loop and the Periplasmic Domain Allosterically Influence YidC’s Other Functionally Important Regions

YidC’s U-shaped hydrophilic groove, exposed on the cytoplasmic side of the membrane bilayer, is essential for the insertion process^49,50^. The membrane proteins enter the YidC groove through the cytoplasmic side of the membrane bilayer during the insertion process. The YidC groove area is filled with water to provide a smooth sliding motion for the entering protein. As a membrane protein advances through the insertion processes, the groove water molecules are expelled from the TM groove. These two variables produce a change in the region’s hydrophobicity, making it more vulnerable to membrane insertion^20,35,81^.

To analyze the water content of the groove inside the TM region, the number of water molecules within the groove region of the YidC protein was quantified and plotted against simulation time. The lack of the C2 loop significantly impacted the amount of water inside the groove area (Fig. 6B & D). Because hydrophilic contacts initially serve to keep membrane proteins in place inside the groove while the insertion process is being carried out^20^. This provides strong evidence that the C2 loop not only contributes to the conformational dynamics of the protein but also plays a significant role in the protein’s function. The water quantity is significantly reduced in system YidC ΔC2 ΔPD loop compared to system YidC. Our findings lead us to infer that the removal of just PD does have a marginal impact on the conformational dynamics of YidC. However, when this modification is coupled with the removal of the C2 loop, the effect is significantly amplified, which could ultimately affect the insertion process.

**Fig. 6.**
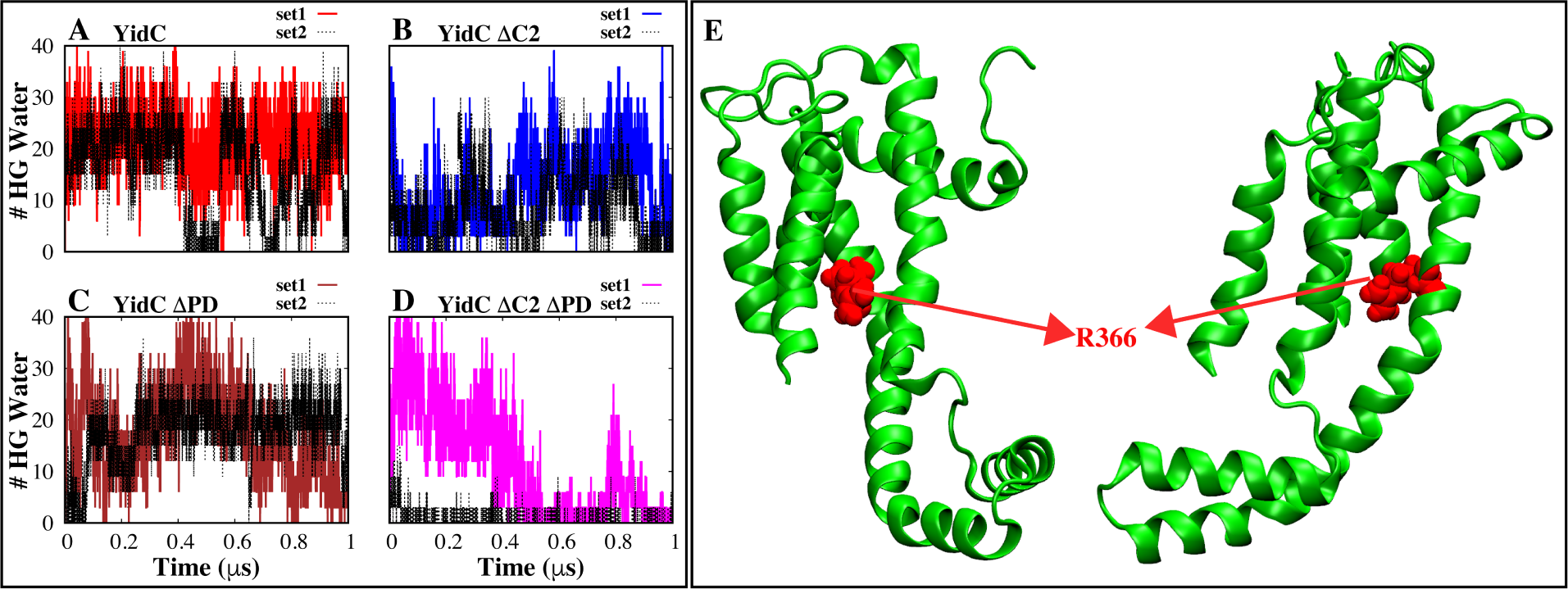
Analyses of the water inside the YidC groove. (A–D) The number of water molecules located within 5 Å of the R366 residue inside the hydrophilic groove (HG) region of YidC in each system. (E) The graphical illustration of the residue R366, which may be found in the central part of the hydrophilic groove (HG) of YidC.

We also found an intra-domain hydrogen bond in the TM region between Y516 and G429 that is only stable in the wild-type system compared to other systems (Fig. 7). Especially in the YidC ΔC2 ΔPD system, this bond is completely broken. We think this hydrogen bond is unstable in the system YidC ΔC2 ΔPD because the fluctuation of the TM region is caused by the absence of C2 loop (Fig 7B & D). These results clearly show a link between the functionally important C2 loop and the PD region on the TM side of YidC. Furthermore, the presence of this hydrogen bond (Fig. 7) appears to play a crucial role in the insertion process. As the incoming membrane protein moves along the groove and toward the periplasmic side, it breaks this link (Fig 7), which causes widening of the TM region and leads the water to leave the hydrophilic groove, resulting in a hydrophobic shift and increasing the likelihood of membrane insertion.

**Fig. 7.**
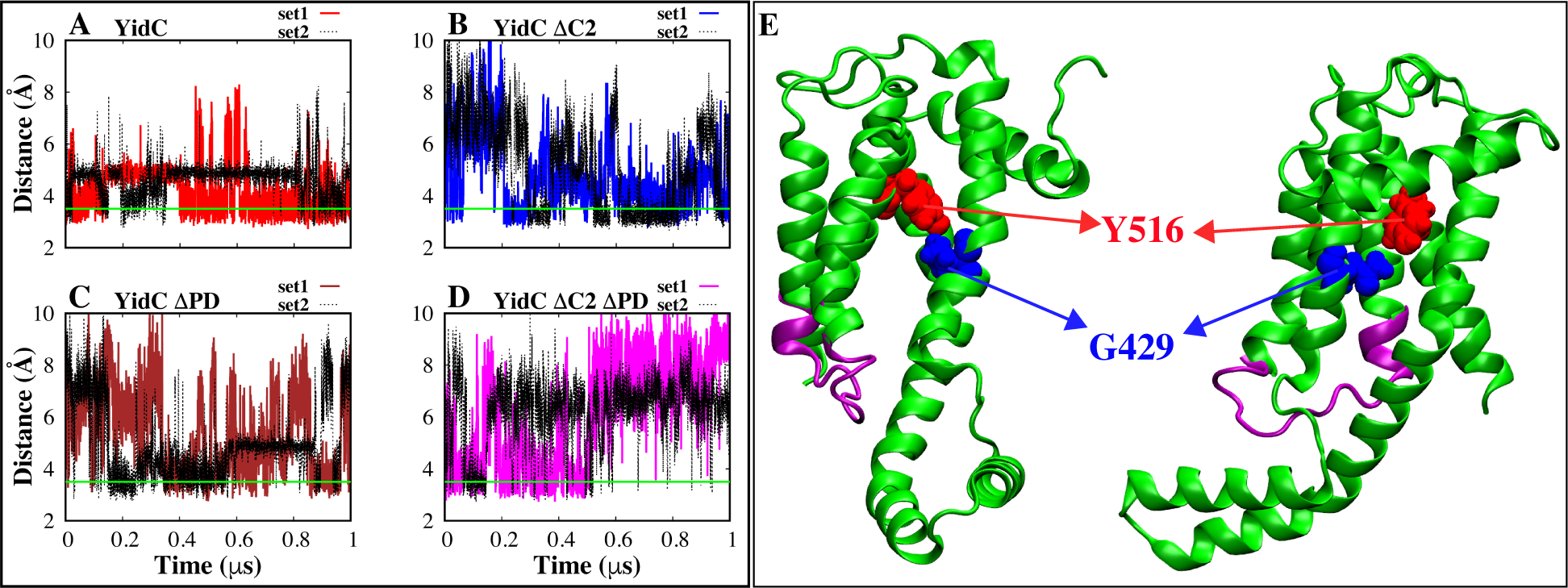
Interaction analysis of a hydrogen bond between Y516 and G429 (YidC), which is situated inside the groove area. (A–D) Analysis of the distance between the hydrogen bonds formed by Y516 and G429. (E) The graphical representation of the residues that participate in the interaction involving hydrogen bonds.

To this point, we have shown that the C2 loop and PD directly alter the structural behavior of YidC, while the PD region also slightly affects the conformational dynamics of the protein. However, the absence of both the PD and the C2 loop has an allosteric influence on the behavior of the YidC conformational dynamics, although the effect of the C2 loop absence is substantially more significant than the PD absence. This leads us to the conclusion that the C2 loop is much more critical to the structure and function of YidC than the PD region, which is in support of previous research^44,45^. On the other hand, removing just PD affected the TM region’s conformational dynamics. To determine what caused this effect, we analyzed interactions between the PD and the TM region that play a significant role in the stability of the protein.

### Inter-domain Amino Acids Interactions Play a Key Role in the Stabilization of the Transmembrane Domain

Previous experimental findings lead researchers to hypothesize that the interactions between the PD region of YidC and the TM region of YidC are crucial for maintaining the protein’s stable state in the membrane^44^. Through this research, we were able to identify critical hydrogen bonds and salt-bridle interactions that take place between the PD region and the TM region. It is interesting to note that the hydrogen bonds formed between the PD region and the TM region were only observed in the wild-type YidC system and entirely disrupted in the YidC ΔC2 system. However, there is no direct interaction between the PD area and the C2 loop region, which does affect the TM region’s stability. We also identified a salt bridge interaction between PD and TM, which is contributing to the stability of the TM region. The salt bridge between K232 and D329 (Fig. 8) is stably formed in the wild-type YidC system; however, this salt bridge is disrupted in YidC ΔC2 (Fig. 8B).

**Fig. 8.**
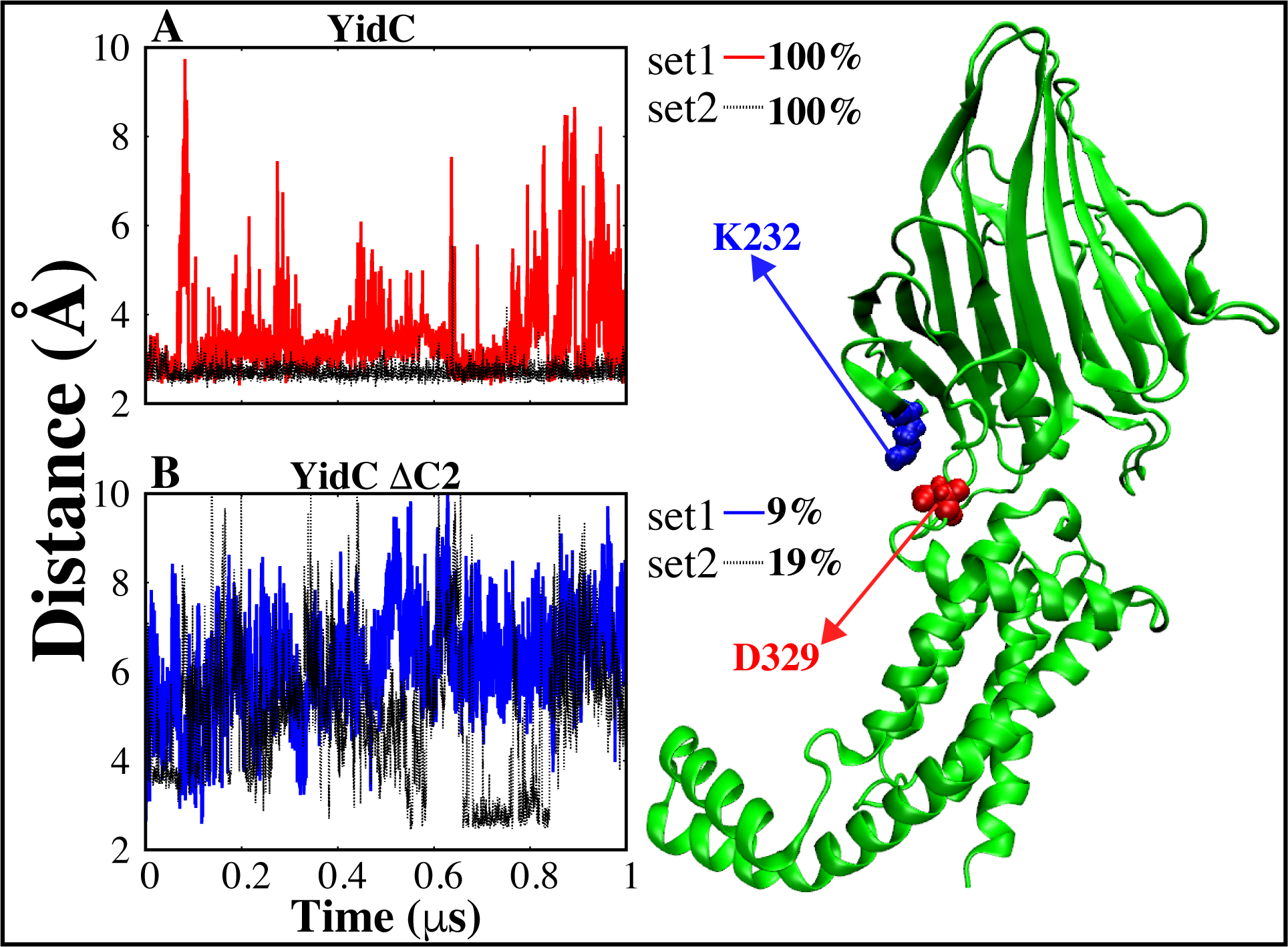
Salt–bridge formed between D329 and K232 (YidC), which is located between the PD and TM regions. The cartoon representation of the salt-bridge interactions that take place between the PD region and the TM area (right). (A–B) Distance analysis between D32 and K232 salt-bridge and we also reported the occupancy of salt-bridge interaction.

**Table 1:** Occupancy (%) of Inter-domain H-Bonds between PD and TM region.

|  | **Y259-M441 (%)** |  | **A214-Y437 (%)** |  | **T327-E312 (%)** |  |
| --- | --- | --- | --- | --- | --- | --- |
| System | set1 | set2 | set1 | set2 | set1 | set2 |
| **YidC** | 40 | 40 | 66 | 42 | 81 | 43 |
| **YidC ΔC2** | 0 | 0 | 0 | 0 | 0 | 0 |
| **YidC ΔPD** | NA | NA | NA | NA | NA | NA |
| **YidC ΔC2 ΔPD** | NA | NA | NA | NA | NA | NA |

We believe that the removal of the C2 loop has caused an effect on the hydrogen bond (Fig. 7) in the TM core region, causing instability of the TM region and ultimately causing disruption of hydrogen bonds and salt bridge interactions between the PD and the TM region. Based on the analysis presented above, we postulate that the hydrogen bond (Fig. 7) in the protein’s groove region is essential for preserving the structural stability of the protein. The C2 loop is crucial for this hydrogen bond to remain stable in the structure. Although the YidC PD region does have a more significant influence on the protein’s structure, its contribution to the protein’s overall function is noticeably less substantial than that of the C2 loop. Even without the PD region, it is still possible to have a normal Sec-independent insertase mechanism; however, the absence of the C2 loop may have a detrimental influence on the protein function.

### Proposed Independent Insertion Mechanism of Gram-negative Bacterial YidC

According to the findings and earlier hypotheses, gram-negative bacterial YidC must likewise undergo significant conformational changes during the Sec-independent insertion procedure, much like gram-positive bacterial YidC. During the Sec-independent insertion process, the entering membrane protein would first contact the cytoplasmic loops and then move into the hydrophilic groove, where it would join forces with R366 to create a salt bridge. Incoming protein must be moved into YidC’s hydrophilic groove by the YidC loops on the cytoplasmic side of the bilayer. The salt bridge between the incoming protein and R366 of YidC also contributes to the passage of the protein towards the periplasmic side, stabilizing its position within the groove. As the incoming protein moves through the groove and approaches the periplasmic side, it breaks the hydrogen bond between Y516 and G429, causing a widening of the TM region that results in a hydrophobic shift through dehydration of the groove. The protein will finally make contact with lipid tails. After that, with the aid of the proton motive force and the membrane’s hydrophobic interaction, the protein moves in the direction of the membrane. Then, via the groove, the protein enters the membrane.

## Conclusions

This work shows that the C2 cytoplasmic loop of YidC could be an essential part of the protein. This could be achieved, for example, by the C2 cytoplasmic loop of YidC helping to stabilize the protein structure via its indirect effects on interactions in the transmembrane core region, as well as its indirect effects on other periplasmic domains, in particular on interactions between the PD and the TM region. It also indicates that the existence of the C2 loop influences the functional features of YidC, such as the hydration of the groove. More studies are required to further understand the role of the C2 loop in the sec-independent insertion process of small single-spanning membrane proteins like the pf3 coat protein, whose interactions with the cytoplasmic region are thought to be essential.

In the context of molecular dynamics simulations, the results of our research show that C2 loops may be required for the structural stability and Sec-independent function of the gram-negative YidC membrane protein dynamics.

## Supporting Information Description

Movie S1 and Figures S1–S3 in Supporting Information provide additional analysis based on our MD simulations as discussed in the manuscript.

Simulation and analysis scripts are available on GitHub Page: https://github.com/bslgroup/YidC.

## Acknowledgement

This research is supported by the National Science Foundation grant CHE 1945465, the National Institutes of Health Grant R15GM139140, R35GM147423, and the Arkansas Biosciences Institute. This work also used the Extreme Science and Engineering Discovery Environment (XSEDE), which is supported by National Science Foundation Grant ACI-1548562. This work used XSEDE resource Stampede through allocation MCB150129. Anton 2 computer time was provided by the Pittsburgh Supercomputing Center (PSC) through Grant R01GM116961 from the National Institutes of Health. The Anton 2 machine at PSC was generously made available by D.E. Shaw Research.

---

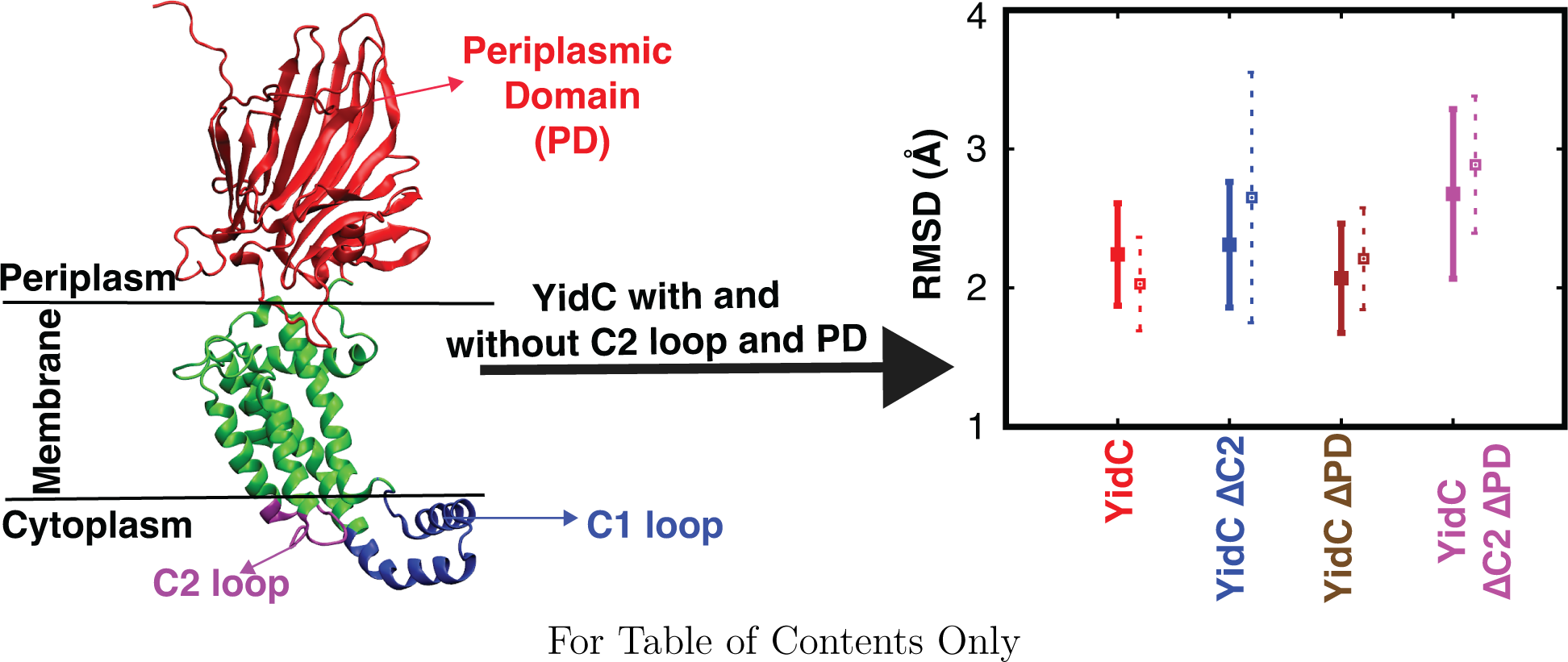
Table of Contents Image.

