## Supporting Information for "Deciphering the Inter-domain Coupling in a Gram-negative Bacterial Membrane Insertase"

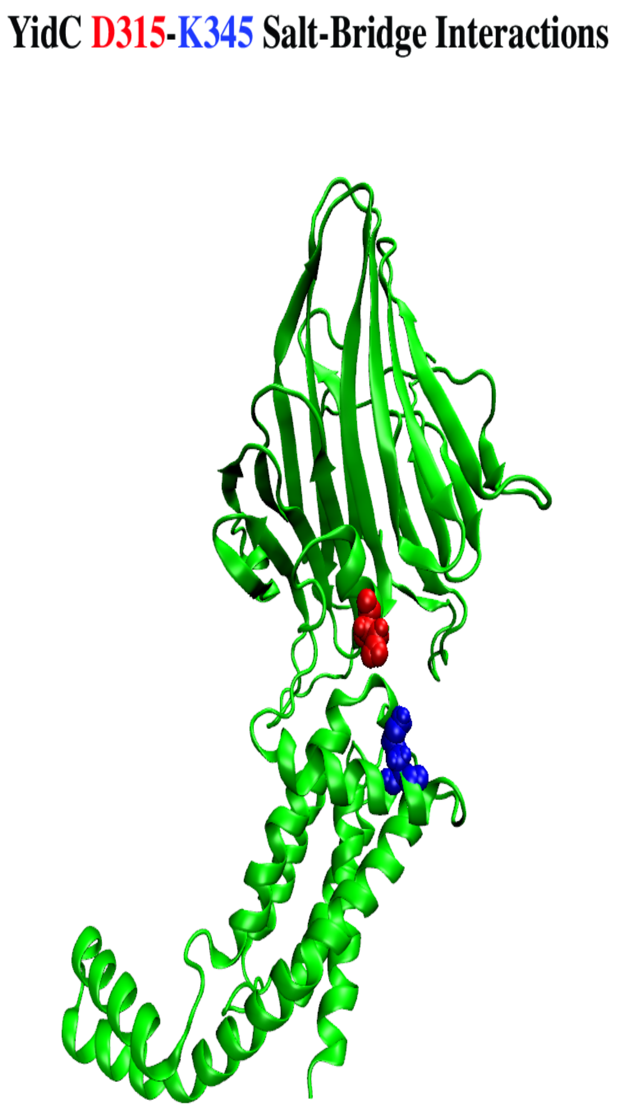


**Mov. S1.** Salt–bridge interaction between D315 and K345 (YidC), which is located between the PD and TM regions.

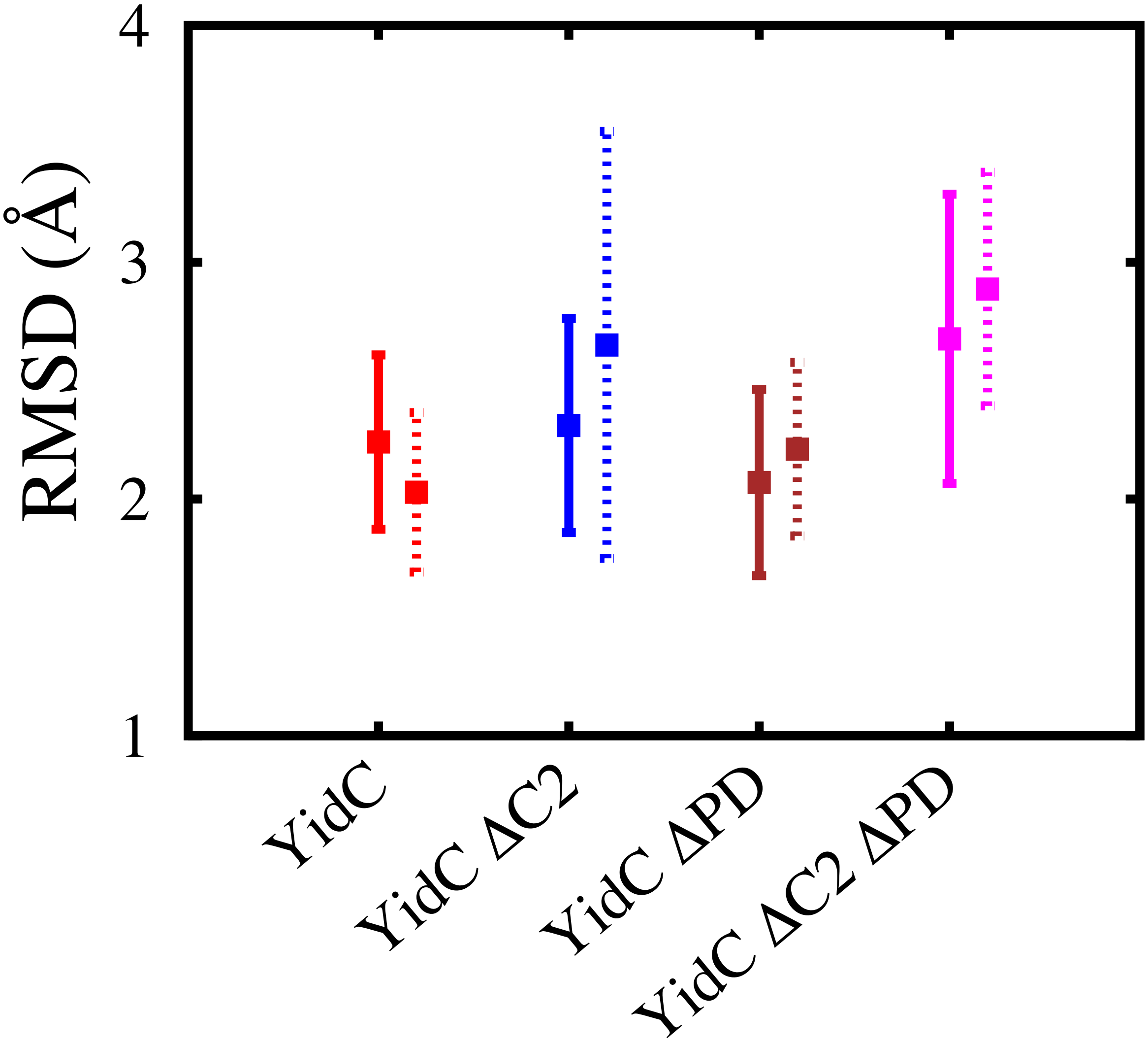


**Fig. S1.** Analysis of the structural stability of YidC with and without the PD and C2 loop. The average and standard deviation of the YidC's root mean square deviation in various systems are shown in this figure. Based on RMSD data, we have shown that YidC is more fluctuating in the system without the C2 loop and PD than in the system with the C2 loop and PD. The dashed lines in the graphs represent the second simulation run for each individual system.

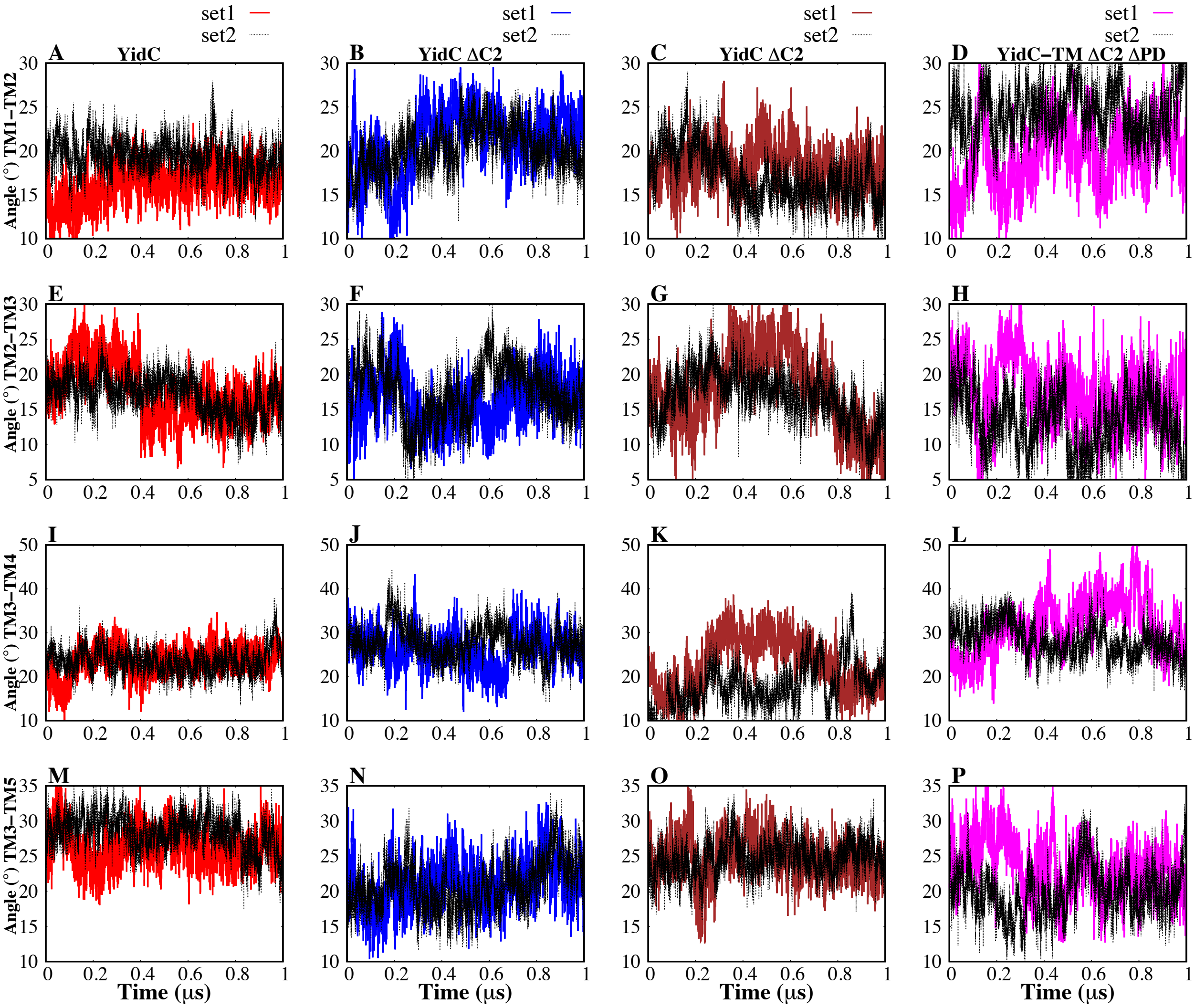


**Fig. S2.** Inter-helical angles between transmembrane helices of YidC. (A–D) The inter-helical angle between the transmembrane helix 1 and helix 2 of the protein. (E–H) The overall inter-helical angle between helix 2 region and helix 3 of the protein. (I–L) The inter-helical angle between the transmembrane helix 3 and helix 4 of the protein. (M–P) The overall inter-helical angle between helix 3 region and helix 5 of the protein.

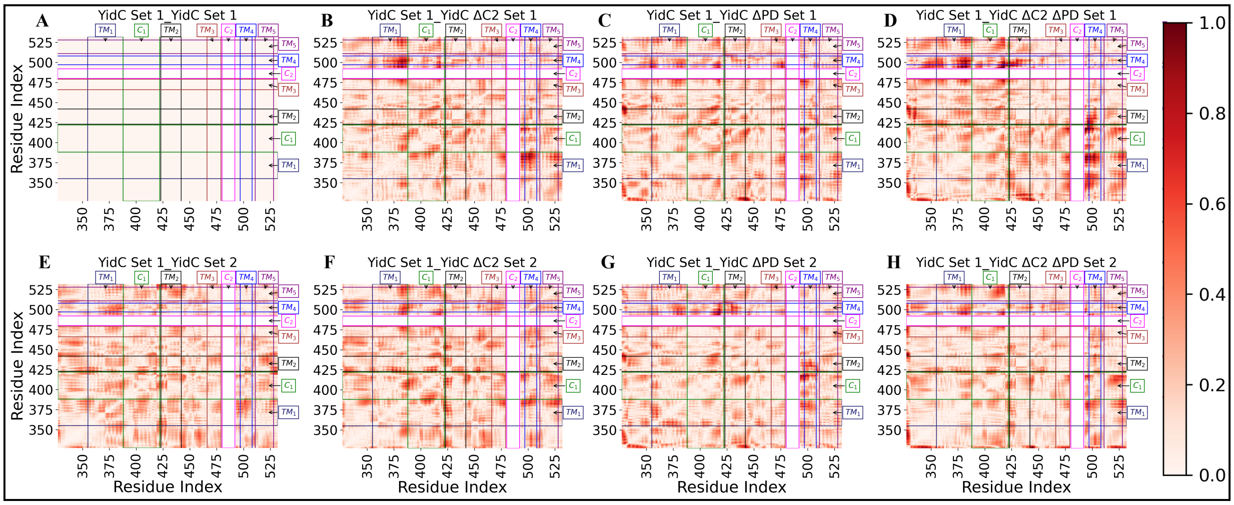


**Fig. S3.** The DNA analysis showed that the YidC set 1 control system and the other YidC systems that were investigated for this research have distinct disparities in their correlations. Although the highest difference in correlation that may be detected in practice was less than one, the theoretical maximum is two. (A-H) Differences in correlation are shown as a gradient of red, with deeper red representing bigger differences.
